# easyEWAS: a flexible and user-friendly R package for Epigenome-Wide Association Study

**DOI:** 10.1101/2025.01.09.632273

**Authors:** Yuting Wang, Meijie Jiang, Siyuan Niu, Xu Gao

**Author notes:** **Corresponding author:** Xu Gao, Ph.D., Department of Occupational and Environmental Health Sciences, School of Public Health, Peking University, No.38 Xueyuan Road, Haidian District, Beijing, China 100191.

## Abstract

**Motivation:** Rapid advancements in high-throughput sequencing technologies especially the Illumina DNA methylation Beadchip greatly fueled the surge in epigenome-wide association study (EWAS), providing crucial insights into intrinsic DNA methylation modifications associated with environmental exposure, diseases, and health traits. However, current tools are complex and less user-friendly to accommodate appropriate EWAS designs and make downstream analyses and result interpretations complicated, especially for clinicians and public health professionals with limited bioinformatic skills.

**Results:** We integrated the current state-of-the-art EWAS analysis methods and tools to develop a flexible and user-friendly R package *easyEWAS* for conducting DNA methylation-based research using Illumina DNA methylation Beadchips. With *easyEWAS*, we provide a battery of statistical methods to support differential methylation position analysis across various scenarios, as well as differential methylation region analysis based on the DMRcate method. To facilitate result interpretation, we provide comprehensive functional annotation and result visualization functionalities. Additionally, a bootstrap-based internal validation was incorporated into *easyEWAS* to ensure the robustness of EWAS results. Evaluation in asthma patients as the example demonstrated that *easyEWAS* could simplify and streamline the conduction of EWAS and corresponding downstream analyses, thus effectively advancing DNA methylation research in public health and clinical settings.

**Availability and implementation:** *easyEWAS* is implemented as an R package and is available at https://github.com/ytwangZero/easyEWAS.

## Introduction

Epigenetics, characterized by modifications to genomic DNA that do not alter the DNA sequence, serves as a bridge linking genetics and environment, thereby providing insights into the pathophysiology of diseases (Bollati and Baccarelli, 2010; Gao, 2024). DNA methylation is one of the most widely studied epigenetic mechanisms, wherein methyl groups are added to specific sites within DNA molecules to regulate gene expression (Mattei *et al*., 2022). Over the past few decades, rapid advancements in high-throughput measurement technologies, especially the Illumina Infinium HumanMethylation BeadChips, have enabled the feasible study of methylation at a whole-genome level, further facilitating the widespread application of epigenome-wide association study (EWAS), which elucidates the associations of DNA methylation changes with environmental exposures, diseases, and other health outcomes (Campagna *et al*., 2021). With the increase in data dimensionality and analytical complexity, there is a growing demand for an updated and efficient tool to conduct EWAS analyses for researchers with a broader interest in clinical medicine and public health.

To date, several bioinformatics tools have been developed to analyze and interpret the high-dimensional methylation data in EWAS, including R packages like *minfi* and *ChAMP*, which primarily utilize ordinary least squares to assess differences in DNA methylation between two groups (Aryee *et al*., 2014; Morris *et al*., 2014). However, the availability of extensive EWAS data allows for research designs beyond simple case-control studies, encompassing more complex designs including cohort studies, randomized crossover trials, time-to-event analyses, and longitudinal studies with repeated measures which are more widely conducted in public health and clinical studies.

Furthermore, these existing tools could address the downstream processing of EWAS results but with challengeable coding, particularly in terms of validation and visualization, which play crucial roles in result interpretation. These obstacles and the steep learning curve for novice researchers and clinicians with limited knowledge of R programming in the field of DNA methylation need to be addressed To consider the aforementioned issues and assist in performing appropriate EWAS analyses in public health and clinical settings, we developed an open-source and user-friendly R package named *easyEWAS* to conduct EWAS efficiently. We then demonstrated an application of *easyEWAS* to explore DNA methylation dynamics in asthma patients using a Gene Expression Omnibus (GEO) database.

## Methods

### Design

The *easyEWAS* tool was primarily developed in the R environment. Tailored specifically for the currently most cited Illumina high-density microarray for genome-wide methylation profiling (Xu and Taylor, 2021), *easyEWAS* aims to simplify and streamline the execution of EWAS and its downstream analyses. Figure 1 illustrates the workflow of the *easyEWAS*, which includes data preprocessing, differential methylation analysis, bootstrap-based internal validation, result visualization, and enrichment analysis.

**Figure 1.**
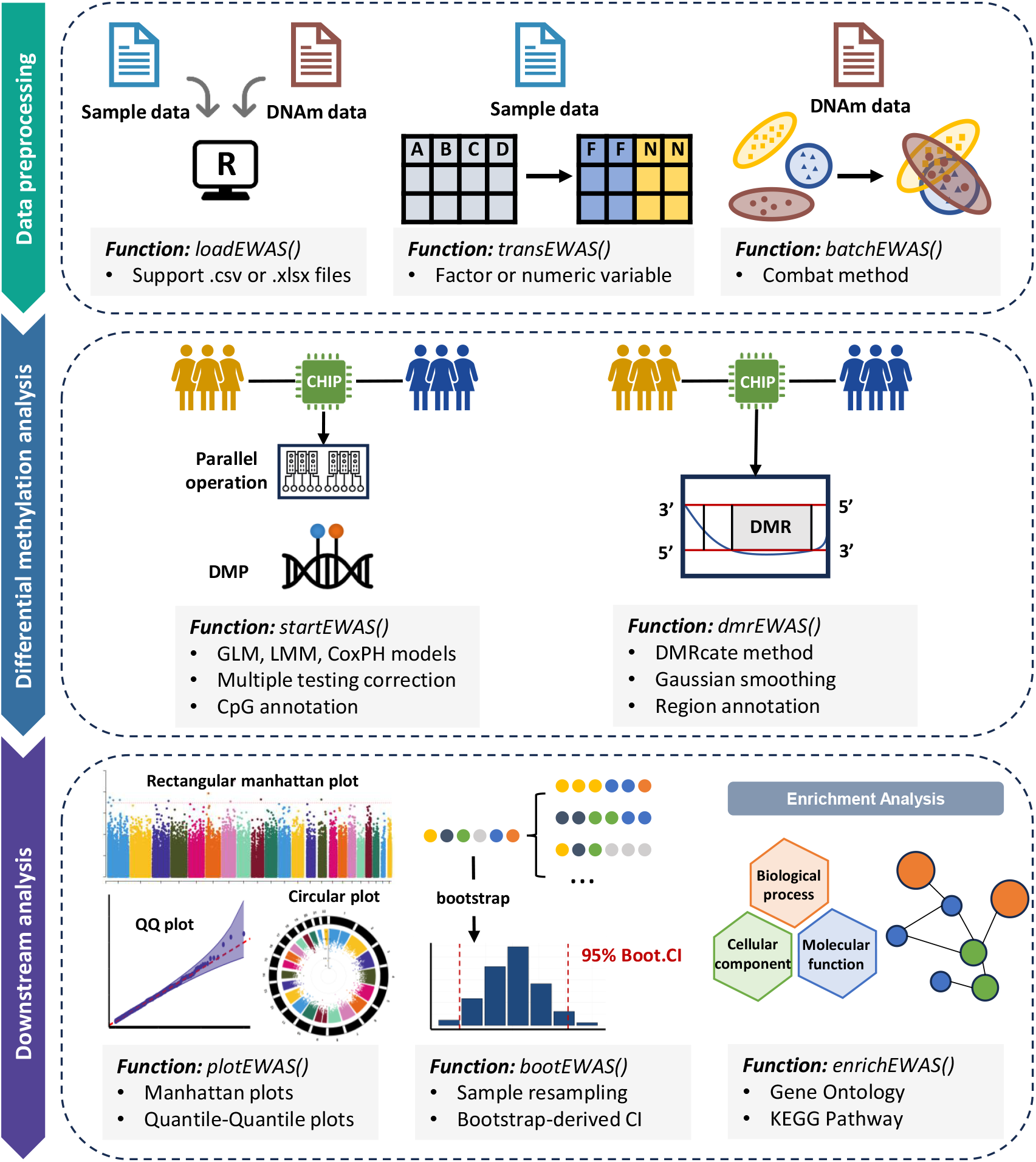
The workflow chart of the *easyEWAS*. **Abbreviations:** DNAm, DNA methylation; DMP, Differentially Methylated Position; DMR, Differentially Methylated Region; GLM, General Linear Model; LMM, Linear Mixed-Effects Model; CoxPH, Cox Proportional Hazards Model; QQ: Quantile-Quantile; Boot.CI, Bootstrap-derived CI; KEGG, Kyoto Encyclopedia of Genes and Genomes.

### Data preprocessing and batch effect correction

The *easyEWAS* package requires users to provide a dataset containing sample information (e.g., the phenotype of interest or other covariates), along with a corresponding dataset of DNA methylation values (either β-values or M-values), which are typically provided by vendors or contractors, or obtained by processing raw IDAT files using specialized packages. It is recommended that users utilize *easyEWAS* to convert variables in sample data to appropriate types (e.g., factor or numeric) to meet the analysis requirements. Considering the potential for batch effects to completely undermine biological results, *easyEWAS* supports adjustment for known batches using an empirical Bayesian framework based on the built-in *sva* package (Leek *et al*., 2012). The output is a set of corrected DNA methylation values with batch effects effectively removed.

### Differential methylation analysis

Differentially Methylated Positions (DMPs) are CpG sites with statistically significant differences with respect to the primary exposure or health outcomes. The DMP analysis is thus the primary differential analysis in EWAS, aimed at identifying positions in the genome where methylation levels exhibit significant changes across different physiological states, disease statuses, tissue types, or environmental conditions (Campagna *et al*., 2021). In the context of common scenarios encountered in EWAS analysis, *easyEWAS* provides three distinct DMP analysis methods (Supplement methods):

- General Linear Model (GLM): Using the methylation level (β value or M value) of each CpG site as the dependent variable, and the phenotype of interest as the independent variable, while simultaneously adjusting for necessary covariates (Formula 1, details in supplement methods). It is commonly employed to compare DNA methylation differences between binary variables (e.g., case-control studies) or multiple groups (e.g., never smokers, former smokers, current smokers), or to assess associations between methylation patterns and continuous variables such as air pollution exposure, etc.

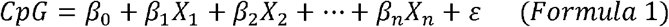

- Linear Mixed-Effects Model (LMM): Unlike GLM, an additional random effects term E is incorporated to capture the random variation in the LMM model (Formula 2). It is typically utilized for analyzing longitudinal studies containing methylation data with two or more time points, incorporating individual variability, between-group variability, or time as random effect terms.

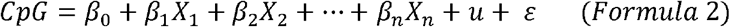

- Cox Proportional Hazards Model (CoxPH): With the methylation level of each CpG site and other covariates serving as the exponential components of the hazard, the model investigates the risk of event occurrence at time t, unveiling the association between DNA methylation patterns and survival time or event incidence (Formula 3).

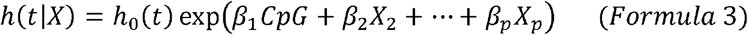

In *easyEWAS*, both GLM and LMM return the regression coefficient for the phenotype of interest, indicating the change in DNA methylation levels when the phenotype changes per one unit (interquartile range, or standard deviation). The output also includes the standard error and an unadjusted *p*-value corresponding to the coefficient. For the CoxPH model, *easyEWAS* provides the hazard ratio (HR) for each CpG site, elucidating the impact of one unit change in methylation level on the risk of event occurrence, along with the corresponding 95% confidence interval (CI) and an unadjusted *p*-value. Necessary annotation information for downstream analysis and result visualization, including chromosome name, coordinate, nearest annotated gene, genomic location, and relation to CpG islands, is also provided for each CpG site in the output. Furthermore, due to the large-scale screening of DMPs which often involves repetitive testing of hundreds of thousands of CpG sites, it is imperative to consider statistical Type I errors. The *easyEWAS* provides two widely adopted methods for correcting significance thresholds: the Bonferroni correction and the Benjamini and Hochberg method (BH method) (Mansell *et al*., 2019). Users can flexibly determine the most suitable correction method based on the specific characteristics of their studies. Notably, by leveraging the *foreach* package with the *doParallel* backend, users of *easyEWAS* can accelerate DMP analyses by specifying the number of logical cores to be utilized, enabling automatic execution of parallel operations. Based on benchmark analysis using simulated data, we recommend utilizing 50%-75% of the available cores for optimal parallel computing performance (Details in supplement methods). Table S1 presents a comparison of the runtime for three different statistical models based on simulated data of the same magnitude (Details in supplement methods).

Compared to DMP, differentially methylated regions (DMRs), which consist of several adjacent DMPs, are often more biologically relevant and more likely to be associated with the expression of modified genes (Campagna *et al*., 2021). The *easyEWAS* package integrates the *DMRcate* method to support DMR analysis. The method first conducts a limma-based regression analysis of the methylation levels at each CpG site in relation to the phenotype, which is independent of the DMP analysis provided by *easyEWAS* in the previous step (Ritchie *et al*., 2015). It then applies Gaussian smoothing to average the effects, followed by grouping neighboring CpG sites within a user-specified window (Peters *et al*., 2015). The default window in *easyEWAS* separates DMRs based on a distance of 1000 base pairs between CpG sites, which is the standard recommendation in most EWAS (Spindola *et al*., 2019). DMR analysis in *easyEWAS* is an optional downstream step, allowing users to decide whether to perform it based on the specific requirements of their study design.

### Bootstrap for internal validation

Validation of the preliminary findings of DMP analysis is crucial to ensure its robustness, as EWAS analyses are typically conducted on large-scale datasets, which may be susceptible to various known or unknown confounding factors (Campagna *et al*., 2021). The *easyEWAS* supports a bootstrap-based internal validation method, which is one of the most effective validation techniques, achieved through computationally intensive resampling techniques (Steyerberg *et al*., 2001). This process generates a multitude of resampled datasets, mirroring the characteristics of the original population. By analyzing each resampled dataset independently, interval distribution of statistical measures such as regression coefficients can be derived, offering insights into the stability and reliability of the preliminary findings obtained through DMP analysis. The *easyEWAS* incorporates five different types of two-sided non-parametric confidence interval calculation methods, including the first-order normal approximation, the basic bootstrap interval, the studentized bootstrap interval, the bootstrap percentile interval, and the adjusted bootstrap percentile interval (Davison and Hinkley, 1997). It allows for the computation of interval distributions for regression coefficients or HR of CpG sites selected based on significance thresholds or specified by users. The *easyEWAS* ultimately provides the original statistical values for each CpG site alongside their bootstrap-derived confidence intervals. To assess the robustness of findings, users can evaluate two key aspects: First, for GLM or LME models, robustness is suggested if the bootstrap confidence interval does not include zero, while for CoxPH models, the interval should not include one, indicating consistent non-null associations across resamples. Second, narrower confidence intervals reflect more precise and reliable estimates, whereas wider intervals indicate greater uncertainty and less stable associations. As robustness criteria may vary depending on the study context and data characteristics, users are encouraged to compare confidence intervals with prior research or within their specific analytical framework to make informed evaluations.

### Result visualization

The result visualization function within the *easyEWAS* is underpinned by the integrated R package *CMplot*. The *easyEWAS* automatically generates Manhattan plots (rectangular or circular), Quantile-Quantile plots (QQ plots), and CpG site density plots based on the results of EWAS. Among these, Manhattan plots are primarily utilized to illustrate the associations between DNA methylation patterns and phenotype of interest. QQ plots assess whether the data distribution adheres to theoretical distributions and could capture systematic biases (Voorman *et al*., 2011), and CpG site density plots could depict the distribution of CpG sites across chromosomes. For various Illumina Methylation Assays, annotation information necessary for visualization is sourced from Bioconductor packages, including *IlluminaHumanMethylation27kanno*.*ilmn12*.*hg19* (Illumina HumanMethylation27k, 27K), *IlluminaHumanMethylation450kanno*.*ilmn12*.*hg19* (Infinium HumanMethylation450 Bead array, 450K), *IlluminaHumanMethylationEPICanno*.*ilm10b4*.*hg19* (Infinium HumanMethylationEPIC Bead array, EPICv1), and *IlluminaHumanMethylationEPICv2anno*.*20a1*.*hg38* (Infinium HumanMethylationEPIC v2 Bead array, EPICv2), respectively. Additionally, *easyEWAS* also supports the annotation of the Infinium Methylation Screening Array (MSA), which was newly released by Illumina in March 2024.

### Enrichment analysis

Enrichment analysis, one of the most widely used techniques for gene list analysis, can facilitate the understanding of the role of DNA methylation in gene regulation and biological processes. The *easyEWAS* supports two functional annotation methods (GO and KEGG) for differentially methylated genes associated with DMPs through the built-in *clusterProfiler* package (Wu *et al*., 2021). Specifically, GO (Gene Ontology, https://www.geneontology.org/) enrichment analysis describes the potential functions of genes in terms of cellular components, molecular functions, and biological processes, thereby inferring related biological mechanisms or disease processes. KEGG enrichment analysis involves comparing the gene list obtained from EWAS analysis with metabolic pathways, cellular processes, and disease information in the Kyoto Encyclopedia of Genes and Genomes database (https://www.genome.jp/kegg/) to reveal significantly enriched biological processes or pathways under specific biological conditions. Moreover, *easyEWAS* also provides visualization functions for enrichment analysis results, including bubble and bar plots.

### Example

To evaluate the performance of the *easyEWAS* package, we downloaded the genome-wide DNA methylation data detected by the Infinium HumanMethylation450 Bead array from the GEO database (GSE148000) to conduct EWAS analysis in asthma patients (Groth et al., 2020). Participants in this study were extracted from the German prospective cohort study ALLIANCE, with all providing sputum samples. Our study included nine non-smoking asthma patients and seven non-smoking healthy controls without any history of pulmonary or respiratory infections. Genomic DNA was drawn from the sputum samples, bisulfite-converted, and hybridized to the Illumina Infinium HumanMethylation450 BeadChip, following the standard Illumina workflow. CpG sites located on sex chromosomes and those associated with SNPs were excluded, resulting in 456,513 CpG sites remaining for further analyses. We then applied the *easyEWAS* to complete the entire analysis pipeline, which primarily involved four steps: preparation, differential methylation analysis, result visualization and enrichment analysis. More details can be found in supplement methods.

## Results

### Implementation

The *easyEWAS* is compatible with Microsoft Windows, Linux, and macOS, allowing seamless installation and execution across these systems. Table 1 presents the complete list of function names and their corresponding functionalities within the *easyEWAS* package. The *easyEWAS* pipeline follows a procedural approach, requiring the sequential execution of each function (Figure 1). It supports differential analyses and downstream analyses for all currently available Illumina Methylation Assays, including 27K, 450K, EPICv1, EPICv2, and MSA. The actual runtime varies with the computer hardware performance.

**Table 1.**
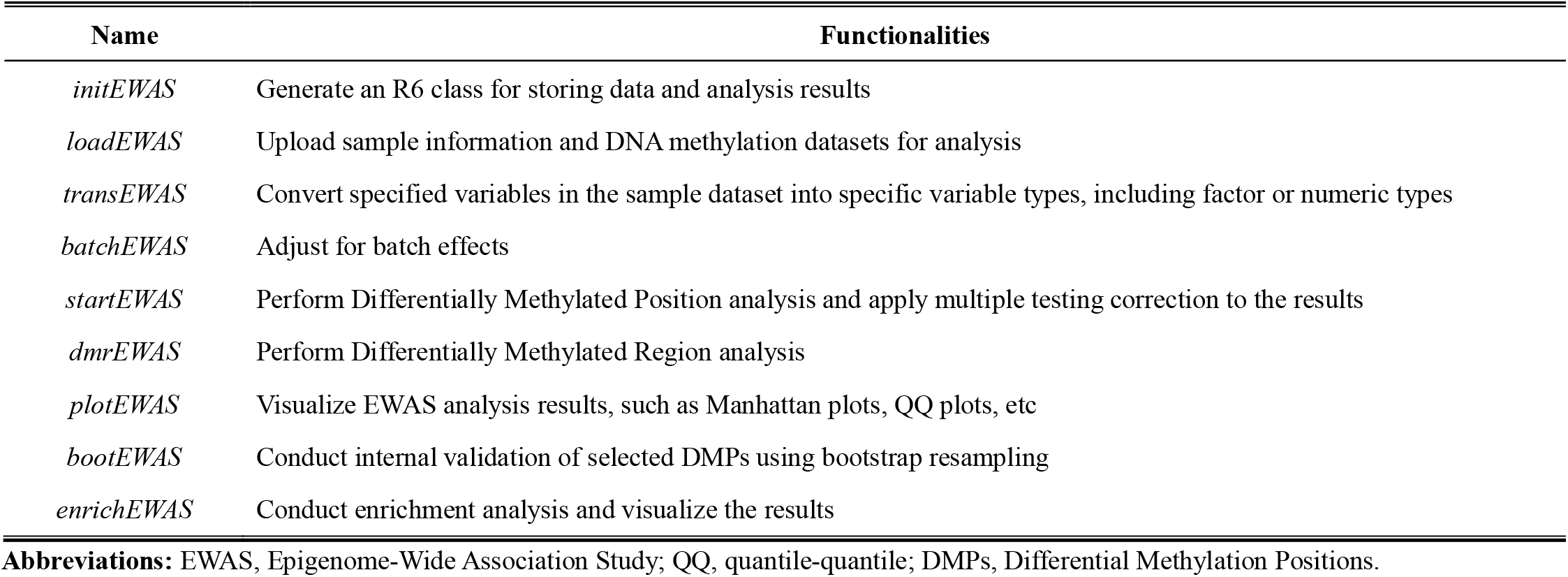
The function names and their corresponding functionalities within the *easyEWAS* package.

### Application in asthma patients

We applied the *easyEWAS* to explore the DNA methylation dynamics between asthma patients and healthy controls. Detailed information and the R code can be found in the supplementary materials. Table S2 provides a summary of the demographic information of nine asthma patients (three females and six males) and seven healthy controls (one females and six males) included in our analysis. The average age of asthma patients was slightly higher than that of healthy controls, but the distribution showed no significant difference. The GLM model was chosen for the EWAS analysis, with age and sex included as covariates. DMP analyses for over 450,000 CpG sites in this study were completed in approximately three minutes using five physical cores on a computer processor with eight physical cores (16 logical cores). Figure S1A illustrate the genome-wide DNA methylation patterns in asthma patients compared to healthy controls. A total of 12 DMPs mapped on six genes across seven chromosomes are presented in Table 2, with Bonferroni-corrected *p*-values < 0.05. Among them, eight CpG sites exhibited significantly lower DNA methylation levels in asthma patients compared to controls. The QQ plot indicates that there may be underlying stratification in the sample that may not have been captured in this study (Figure S1B). We further utilized the *easyEWAS* to perform bootstrap-based internal validation (500 iterations) on the preliminary DMP analysis results. The distribution of confidence intervals for all 12 DMPs confirmed the robustness of our findings (Table S3). Table S4 and Figure S1C presented 12 biological functions with adjusted *p*-values <0.05 in GO enrichment analysis. Table S5 provides the details of 136 DMRs with an FDR < 0.05 detected between asthma patients and healthy controls.

**Table 2.** Characteristics of differential methylation CpG sites between asthmatic patients and healthy controls ^a^.

| probe | Coefficient <sup>b</sup> | SE | *p*-value <sup>b</sup> | FDR | Bonferroni | CHR | Relation to island | Gene | Location |
| --- | --- | --- | --- | --- | --- | --- | --- | --- | --- |
| cg00394712 | 0.405 | 0.013 | <0.001 | <0.001 | <0.001 | 1 | Island | *NUDT17* | TSS200 |
| cg02449698 | -0.128 | 0.011 | <0.001 | 0.003 | 0.028 | 1 | OpenSea | *EFCAB7* | Body |
| cg04529486 | -0.053 | 0.005 | <0.001 | 0.003 | 0.032 | 3 | S_Shelf | / | / |
| cg07996580 | 0.038 | 0.003 | <0.001 | 0.004 | 0.045 | 1 | OpenSea | / | / |
| cg11527367 | -0.081 | 0.007 | <0.001 | 0.003 | 0.026 | 1 | OpenSea | *C1orf152* | Body |
| cg13391740 | 0.052 | 0.004 | <0.001 | 0.002 | 0.006 | 6 | S_Shelf | / | / |
| cg14018420 | 0.139 | 0.011 | <0.001 | 0.003 | 0.015 | 1 | Island | *NBPF3* | TSS1500 |
| cg16282910 | -0.100 | 0.008 | <0.001 | 0.003 | 0.023 | 9 | Island | / | / |
| cg19725474 | -0.029 | 0.002 | <0.001 | 0.003 | 0.024 | 14 | OpenSea | / | / |
| cg23106865 | -0.139 | 0.008 | <0.001 | <0.001 | <0.001 | 7 | OpenSea | / | / |
| cg23835894 | -0.043 | 0.002 | <0.001 | <0.001 | <0.001 | 22 | Island | *GATSL3* | TSS200 |
| cg26670636 | -0.103 | 0.009 | <0.001 | 0.003 | 0.036 | 7 | OpenSea | *C7orf61* | TSS1500 |

**Abbreviations:** CpG, cytosine-guanine dinucleotide; SE, standard error; FDR, false discovery rate; CHR, chromosome.
**a:** Models adjusted for age (years) and sex (male/female).
**b:** The regression coefficient, standard error and *p*-value are derived from the general linear model.
/: unassigned to any genes

### Performance comparison with existing EWAS tools

Table S6 provides a comprehensive comparison of the functionalities offered by *easyEWAS, minfi* and *ChAMP* for EWAS pipeline analysis starting from preprocessed β- or M-values. Unlike *minfi* and *ChAMP*, which partially rely on third-party packages for DNA methylation array support and annotation, *easyEWAS* natively supports all major Illumina Methylation Assays, including the latest MSA, and integrates direct annotation of CpG sites. A key strength of *easyEWAS* lies in its enhanced functionality for DMP analysis, offering a suite of statistical models to accommodate a range of study designs, along with parallel computing capabilities to improve efficiency. Furthermore, *easyEWAS* incorporates features absent in the other tools, including bootstrap-based internal validation and advanced visualization options, to enhance the results interpretations. By addressing critical gaps in existing tools, *easyEWAS* provides a more efficient solution for epigenetic epidemiology research.

## Discussion

*easyEWAS* is a flexible and user-friendly R package that systematically performs EWAS analyses under various study designs, along with downstream analyses and results visualization. During the development of *easyEWAS*, careful consideration was given to selecting appropriate methods for each analysis module. For instance, we selected the *ComBat* method for batch effect correction due to its robustness with small sample sizes, its flexibility in allowing users to retain covariates, and its suitability for DNA methylation data with known batch or plate effects (Leek *et al*., 2012). For DMR analysis, we selected *DMRcate* method due to its ability to aggregate adjacent CpG sites using Gaussian kernel smoothing, reducing noise from irregular spacing. Unlike methods reliant on genomic annotations, *DMRcate* defines DMRs based on CpG spatial coordinates, enabling the detection of regulatory regions beyond standard annotations. It has demonstrated superior accuracy and sensitivity, particularly for small or complex effects, and its user-friendly interface aligns well with the design principles of *easyEWAS* (Peters *et al*., 2015, 2021).

Compared to existing tools, our package offers several unique advantages. First, it boasts fast execution speed and simple operation, requiring minimal code to automate a complete EWAS analysis, making it particularly suitable for beginners in the field of epigenetic epidemiology. Second, the package incorporates multiple statistical models, catering to a wide range of study designs in current EWAS analyses beyond traditional case-control studies. Ultimately, it streamlines the EWAS and its downstream analyses, providing a comprehensive analytical pipeline for epigenetic research. Despite these strengths, *easyEWAS* has certain limitations that should be acknowledged. First, it supports pipeline analysis only for preprocessed DNA methylation data (β-values or M-values) and does not handle raw IDAT files. However, in most clinical or research settings, vendors or contractors typically provide preprocessed DNA methylation data. Moreover, several well-established packages already exist for processing raw IDAT files, allowing *easyEWAS* to focus on refining the epigenetic epidemiology analysis pipeline (Aryee *et al*., 2014; Morris *et al*., 2014). Particularly, since the incorporation of WGBS data processing is being considered in future updates, ensuring compatibility and adaptability remains a priority. Second, bootstrap-based internal validation may impose substantial computational costs in large-sample scenarios. Nevertheless, in *easyEWAS*, this validation is restricted to selected significant sites (e.g., FDR < 0.05) or user-specified sites. This targeted approach substantially reduces the computational burden, making the cost manageable while maintaining analytical rigor.

In conclusion, *easyEWAS* is an R package that can be easily integrated into various DNA methylation microarrays detected by Illumina HumanMethylation Bead Chip, significantly enhancing the accessibility of EWAS.

## Dependencies: Packages and Databases

|  |  |
| --- | --- |
| *clusterProfiler* | https://www.bioconductor.org/packages/release/bioc/html/clusterProfiler.html |
| *CMplot* | https://CRAN.R-project.org/package=CMplot |
| *DMRcate* | https://www.bioconductor.org/packages/release/bioc/html/DMRcate.html |
| *IlluminaHumanMethylation27kanno.ilmn12.hg19* | https://bioconductor.org/packages/release/data/annotation/html/IlluminaHumanMethylation27kanno.ilmn12.hg19.html |
| *IlluminaHumanMethylation450kanno.ilmn12.hg19* | https://bioconductor.org/packages/release/data/annotation/html/IlluminaHumanMethylation450kanno.ilmn12.hg19.html |
| *IlluminaHumanMethylationEPICanno.ilm10b4.hg19* | https://bioconductor.org/packages/release/data/annotation/html/IlluminaHumanMethylationEPICanno.ilm10b4.hg19.html |
| *IlluminaHumanMethylationEPICv2anno.20a1.hg38* | https://bioconductor.org/packages/release/data/annotation/html/IlluminaHumanMethylationEPICv2anno.20a1.hg38.html |
| Infinium Methylation Screening Array | https://support.illumina.com.cn/downloads/infinium-methylation-screening-manifest-files.html |
| Gene Expression Omnibus | https://www.ncbi.nlm.nih.gov/geo/ |
| Gene Ontology | https://www.geneontology.org/ |
| Kyoto Encyclopedia of Genes and Genomes | https://www.genome.jp/kegg/ |
| *sva* | https://bioconductor.org/packages/release/bioc/html/sva.html |

## Data Availability Statement

DNA methylation datasets were downloaded from the Gene Expression Omnibus (https://www.ncbi.nlm.nih.gov/geo/query/acc.cgi?acc=GSE148000). The R package *easyEWAS* is available at https://github.com/ytwangZero/easyEWAS, with a tutorial at https://easyewas.github.io/.

