## Supplement Materials for "easyEWAS: a flexible and user-friendly R package for Epigenome-Wide Association Study"

**Table of contents**

**Supplement Methods**

**Table S1** Runtime Comparison of GLM, LMM, and CoxPH Models.

**Table S2** Characteristics of 9 asthmatic patients and 7 healthy controls.

**Table S3** The 95% confidence intervals of regression coefficients based on bootstrap method.

**Table S4** The GO enrichment analysis results of differentially methylated genes corresponding to CpG sites with FDR < 0.05.

**Table S5** Characteristics of differential methylation regions between asthmatic patients and healthy controls.

**Table S6** Comparison of functionality across easyEWAS, minfi, and ChAMP for EWAS pipeline analysis.

**Figure S1** The results obtained from EWAS analyses in asthmatic patients using *easyEWAS*. **(A)** Rectangular Manhattan plot showing *p*-values for epigenome-wide association studies comparing DNA methylation patterns between asthma patients and healthy controls. The red dotted line indicates the *p*-value threshold of the Bonferroni correction method. Abbreviations: Chr, chromosome. **(B)** EWAS quantile-quantile plot showing inflation factor with 95% confidence intervals. **(C)** Bubble plot of GO enrichment analysis of differentially methylated genes.
**Figure S2** Benchmarking analysis of core usage for EWAS runtime optimization.

**Supplement Methods**

***Differentially methylated position analysis***

The three differentially methylated position analysis methods provided by *easyEWAS* include the general linear model (GLM), linear mixed effects model (LMM), and Cox proportional hazards model (CoxPH).

- **General linear model**

For comparing DNA methylation differences between continuous or categorical variables of interest, we use the *lm()* function from *base R*.

$$CpG=\beta_{0}+\beta_{1}X_{1}+\beta_{2}X_{2}+\ldots+\beta_{n}X_{n}+\varepsilon$$

where$CpG$ represents the DNA methylation level for each CpG site, $\beta_{0}$ is the intercept term, $\beta_{1}$is the coefficient associated with the variable of interest$X_{\text{1}}$, and$\beta_{2},\ldots,\beta_{n}$ are the coefficients associated with the covariates $X_{2},X_{3},\ldots,X_{n}$ respectively.$\varepsilon$ denotes the error term accounting for unexplained variability in the methylation level.

- **Linear mixed effects model**

For data with hierarchical structures or repeated measures, we use the *lmer()* function from the *lmerTest* package.

$$CpG=\beta_{0}+\beta_{1}X_{1}+\beta_{2}X_{2}+\ldots+\beta_{n}X_{n}+u+ \varepsilon$$

where $CpG$ represents the DNA methylation level, modeled as a function of fixed effects predictors $X_{1},X_{2},\ldots,X_{n}$, each with respective coefficients$\beta_{1},\beta_{2},\ldots,\beta_{n}$. The term *u* indicates a random intercept term, suggesting variability across different groups or clusters. The error term $\varepsilon$ captures unexplained variability in methylation levels.

- **Cox proportional hazards model**

For survival analysis, we fit a separate CoxPH model for each CpG site using the *coxph()* function from the *survival* package.

$$h\left( t | X \right)=h_{0}\left( t \right)\exp\left( \beta_{1}CpG+\beta_{2}X_{2}+\ldots+\beta_{p}X_{p} \right)$$

where $h\left( t | X \right)$ represents the conditional hazard function at time $t$ given covariates$X$, $h_{0}\left( t \right)$ denotes the baseline hazard function, indicating the hazard at time $t$ when all covariates$X$ are equal to 0. The coefficients $\beta_{1},\beta_{2},\ldots,\beta_{p}$ in the model signify the magnitude of influence of each covariate on the hazard of the event, with$CpG$ representing the methylation level of each CpG site, and $X_{2},X_{3},\ldots,X_{p}$ representing other covariates that may affect the hazard of the event.

***Application in Asthma Patients***

The analysis was performed on a Lenovo device equipped with an Intel® Core™ Ultra 5 125H processor (1.20 GHz), 32.0 GB of RAM (31.6 GB available), and a 64-bit operating system based on x64 architecture.

- **Step 1: Data Preparation**

We first provided the *easyEWAS* package with sample data (“sampledata”) indicating asthma status and covariate information, as well as methylation data (“methydata”) containing β-values representing DNA methylation levels for each sample. Categorical variables such as sex and disease status were converted to factor type to facilitate analysis.

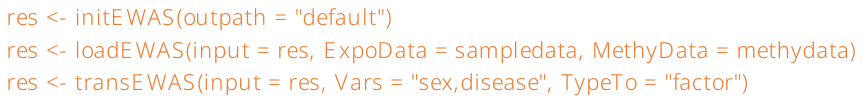

- **Step 2: DMP & DMR Analyses**

The GLM model was chosen for DMP analysis, with age and sex included as covariates. Differential methylation analyses for over 450,000 CpG sites in this study were completed in approximately 3 minutes using five physical cores on a computer processor with 8 physical cores (16 logical cores). We set `adjust = TRUE` to perform multiple testing correction on the results, with Bonferroni-corrected *p*-values < 0.05 identifying sites as differentially methylated between asthma patients and healthy controls.

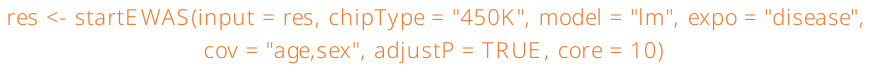

The *dmrEWAS()* function is used to perform DMR analysis between asthma patients and healthy controls, with age and sex adjusted as covariates. Significant CpG sites with gaps of 1000 bp or more will be placed in separate DMRs.

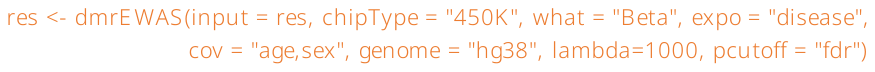

- **Step 3: Result Visualization**

We used the *plotEWAS()* function of *easyEWAS* to create a Manhattan plot, QQ plot, and CpG site density plot of the results using the unadjusted raw *p*-values.

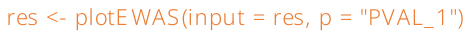

- **Step 4: Internal Validation**

CpG sites with Bonferroni-corrected *p*-values < 0.05 were extracted for bootstrap-based internal validation (500 iterations). The bootstrap percentile interval method was used to calculate the interval distribution of the regression coefficients for each site.

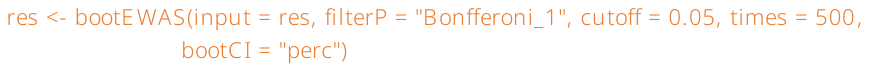

- **Step 5: Enrichment Analysis**

The differentially methylated genes corresponding to CpG sites with FDR < 0.05 were extracted for GO enrichment analysis. A bubble plot was created to visualize the count and gene ratio of each enriched term.

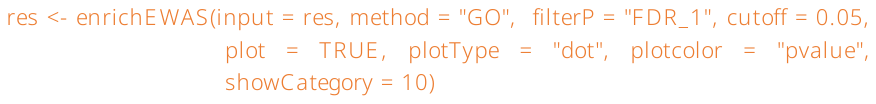

**Benchmark analysis of the optimal number of cores**

To compare the optimal number of cores for parallel computing across a broader range, we conducted the benchmarking on a more powerful server: a Dell PowerEdge T440, equipped with the following specifications: Microsoft Windows Server 2016 Datacenter operating system, dual Intel® Xeon® Silver 4208 CPUs running at 2.10 GHz, and 319.25 GB of installed RAM. We simulated 450K methylation data for 10 individuals and conducted an EWAS analysis based on linear regression (Code available at <https://github.com/ytwangZero/easyEWAS_materials>). We incrementally increased the number of physical cores from five to fifteen and compared the computation times. As shown in Figure S2, computation time decreased significantly with an increasing number of cores within the range of five to nine physical cores. However, beyond nine cores, the overhead associated with task distribution offset the benefits of additional cores. Therefore, we recommend users select 50–75% of their available cores to optimize computational efficiency based on their specific hardware.

**Runtime Comparison of Different Analytical Models**

We generated three sets of 450K DNA methylation data for 10 individuals and compared the runtime of GLM, LMM, and CoxPH models using the *startEWAS()* function in the *easyEWAS* package, with five physical cores. As shown in Table S1, the GLM model was the fastest, completing in 193 seconds, followed by the CoxPH model at 268 seconds. The LMM model was the slowest due to its higher complexity, involving covariance matrix calculations and iterative optimization for random effects, making it significantly more computationally intensive than the CoxPH and GLM models. However, the actual computation time depends heavily on the user’s hardware configuration, including memory and physical cores, so the reported times only serve as general guidance.

**Table S1 Runtime Comparison of GLM, LMM, and CoxPH Models.**

|  | **GLM** | **LMM** | **CoxPH** |
| --- | --- | --- | --- |
| **Sample size** | 10 | 10 | 10 |
| **DNAm array** | 450K | 450K | 450K |
| **Physical cores** | 5 | 5 | 5 |
| **Runtime (seconds)** | 193 | 2323 | 268 |

**Abbreviations:** GLM, General Linear Model; LMM, Linear Mixed-Effects Model; CoxPH, Cox Proportional Hazards Model; DNAm, DNA methylation; 450K, Infinium HumanMethylation450 Bead array.

**Table S2 Characteristics of 9 asthmatic patients and 7 healthy controls ^a^.**

|  | **Asthma**  **(N=9)** | **Healthy**  **(N=7)** | **Overall**  **(N=16)** | ***p*-value ^b^** |
| --- | --- | --- | --- | --- |
| **Sex** |  |  |  |  |
| female | 3 (33.3%) | 1 (14.3%) | 4 (25.0%) | 0.771 |
| male | 6 (66.7%) | 6 (85.7%) | 12 (75.0%) |  |
| **Age (year)** |  |  |  |  |
| Mean (SD) | 58.9 (14.3) | 43.9 (17.2) | 52.3 (16.9) | 0.087 |

**a:** Continuous variables are summarized as mean (standard deviation), and categorical variables as counts (%).

**b:** The *p*-values intervals were calculated using *t* test for continuous variables and Kruskal-Wallis test for categorical variables.

**Abbreviations:** SD, standard deviation.

**Table S3 The 95% confidence intervals of regression coefficients based on bootstrap method.**

| **probe** | **Original Estimate ^a^** | **95% CI lower limit** | **95% CI upper limit** |
| --- | --- | --- | --- |
| cg00394712 | 0.405 | 0.378 | 0.427 |
| cg02449698 | -0.128 | -0.154 | -0.104 |
| cg04529486 | -0.053 | -0.063 | -0.041 |
| cg07996580 | 0.038 | 0.030 | 0.047 |
| cg11527367 | -0.081 | -0.098 | -0.069 |
| cg13391740 | 0.052 | 0.044 | 0.059 |
| cg14018420 | 0.139 | 0.118 | 0.171 |
| cg16282910 | -0.100 | -0.118 | -0.085 |
| cg19725474 | -0.029 | -0.035 | -0.025 |
| cg23106865 | -0.139 | -0.164 | -0.123 |
| cg23835894 | -0.043 | -0.047 | -0.040 |
| cg26670636 | -0.103 | -0.122 | -0.081 |

**a:** The original estimate was derived from the general linear model, adjusting for age (years), sex (male/female).

**b:** The 95% confidence intervals were calculated using the bootstrap percentile method.

**Abbreviations:** CI, confidence interval.

**Table S6 Comparison of functionality across *easyEWAS*, *minfi*, and *ChAMP* for EWAS pipeline analysis.**

| **Feature** | **minfi** | **ChAMP** | **easyEWAS** |
| --- | --- | --- | --- |
| Supported Arrays | 450K, EPIC v1, EPIC v2 (requires third-party annotation packages) | Depends on *minfi* for underlying support | 27K, 450K, EPIC v1, EPIC v2, and MSA |
| DNAm Data Import | Raw .idat files or preprocessed values | Raw .idat files or preprocessed values | Preprocessed β-values, M-values stored in .csv or .xlsx files, or already loaded into the R environment. |
| Data Preprocessing | **×** | **×** | Supports conversion of factor and numeric covariate types in sample data |
| Batch Effect Correction | **×** | ComBat method | ComBat method |
| DMP Analysis | Linear regression and F-test | Limma-based DMP detection | GLM, LMM, and CoxPH models for diverse study designs |
| Results Annotation | **×** | Supports annotation of CpG information | Supports annotation of CpG information |
| Results Visualization | **×** | **×** | Manhattan plots, QQ plots, and CpG density plots |
| Internal Validation | **×** | **×** | Bootstrap-based validation method |
| DMR Analysis | Bump Hunting and Block  Finding methods | Bump Hunting, Probe Lasso, and DMRcate methods | DMRcate method |
| Enrichment Analysis | **×** | Gene Set Enrichment Analysis | GO and KEGG analysis with visualization support |
| Parallel Operation | **×** | **×** | Supported in DMP analysis |

**Abbreviations**: EWAS, Epigenome-wide association study; DNAm, DNA methylation; MSA, Infinium Methylation Screening Array; DMP, Differentially Methylated Position; GLM, General linear model; LMM, Linear mixed effects model; CoxPH, Cox proportional hazards model; QQ plots, Quantile-Quantile plots; DMR, Differentially Methylated Region; GO, Gene Ontology; KEGG, Kyoto Encyclopedia of Genes and Genomes.
