## Supplement Figures for "easyEWAS: a flexible and user-friendly R package for Epigenome-Wide Association Study"

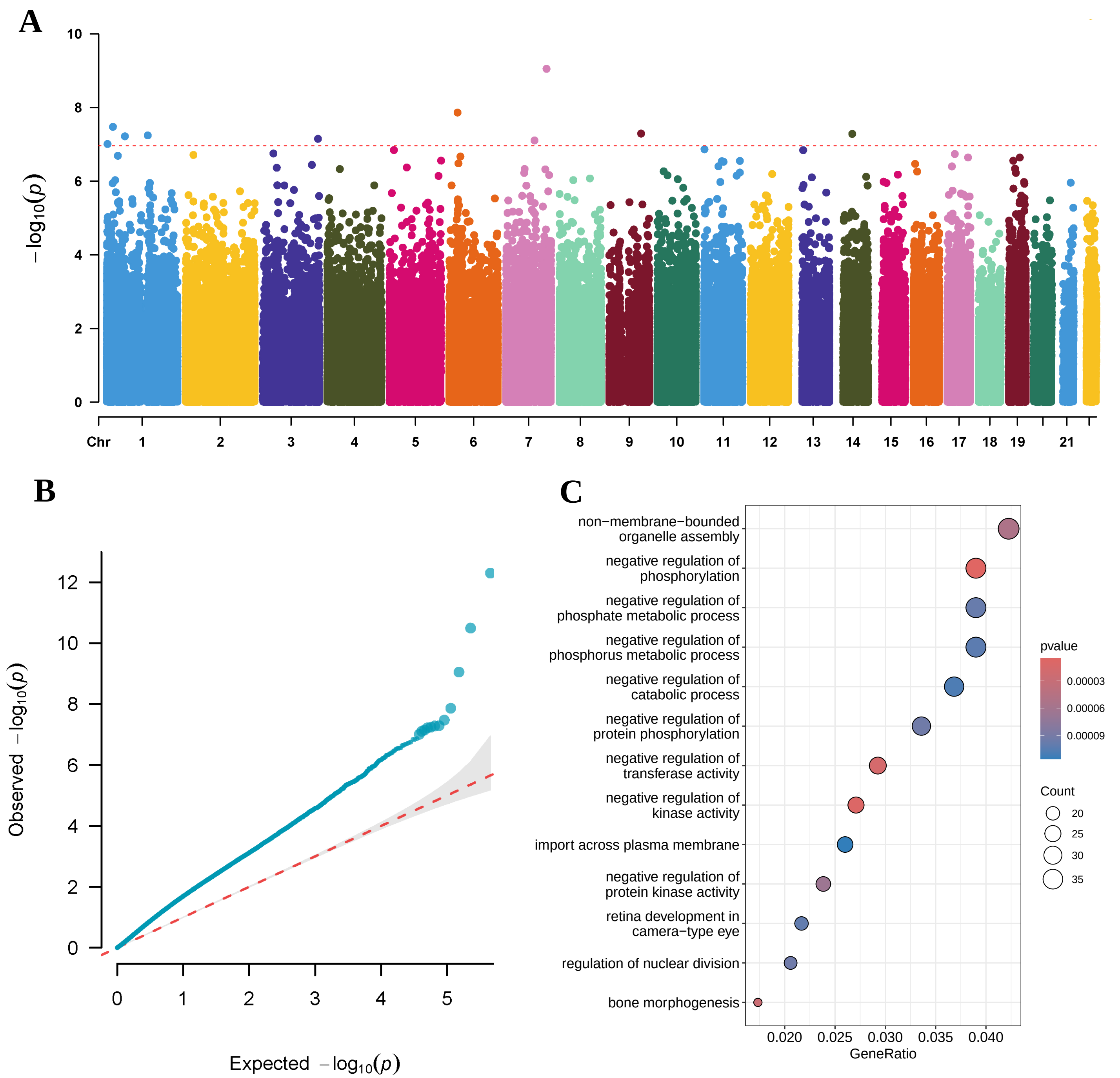


**Figure S1** The results obtained from EWAS analyses in asthmatic patients using *easyEWAS*. **(A)** Rectangular Manhattan plot showing p-values for epigenome-wide association studies comparing DNA methylation patterns between asthma patients and healthy controls. The red dotted line indicates the p-value threshold of the Bonferroni correction method. Abbreviations: Chr, chromosome. **(B)** EWAS quantile-quantile plot showing inflation factor with 95% confidence intervals. **(C)** Bubble plot of GO enrichment analysis of differentially methylated genes.

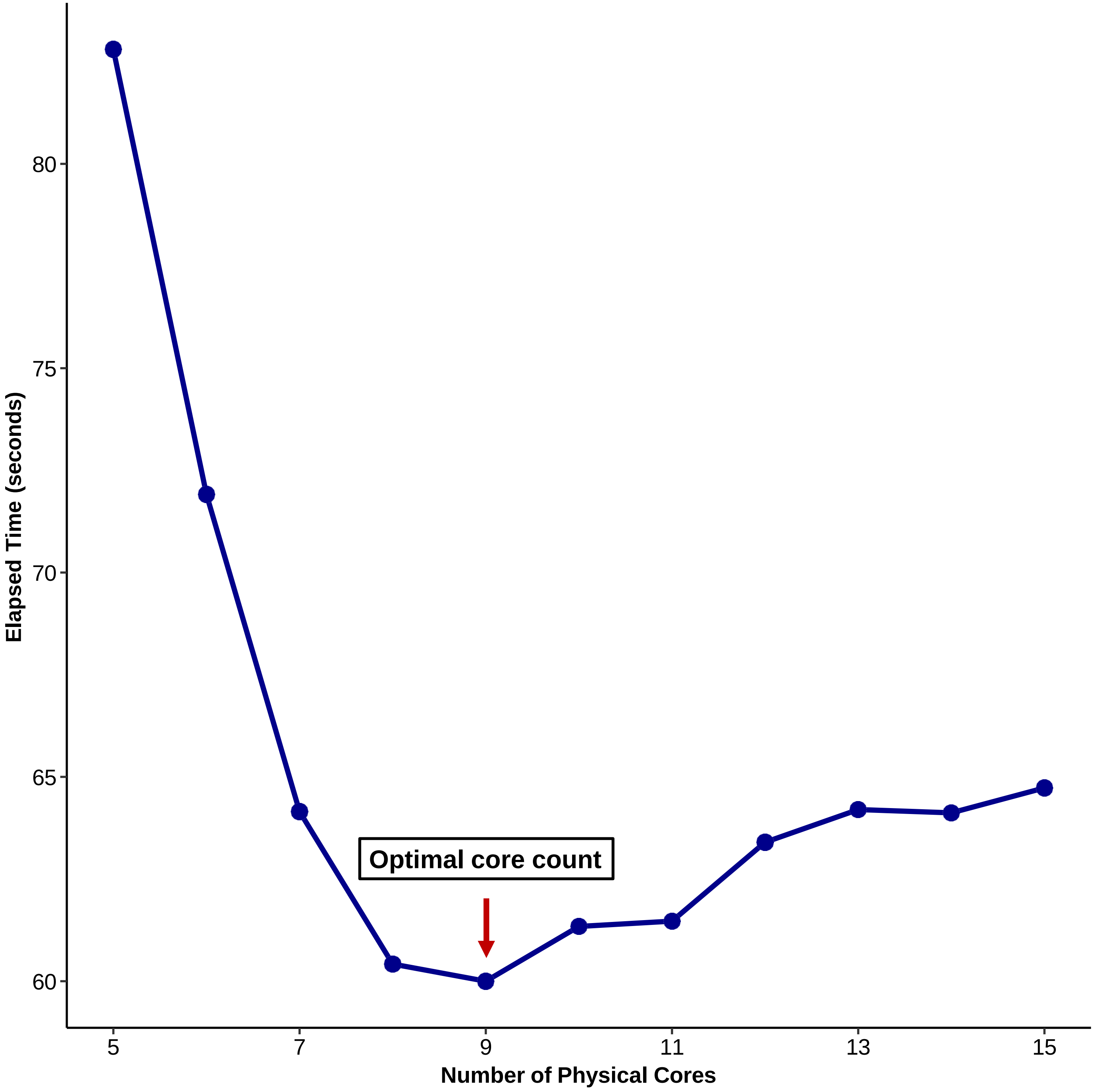


**Figure S2** Benchmarking analysis of core usage for EWAS runtime optimization.
